# Parallel changes in gut microbiome composition and function in parallel local adaptation and speciation

**DOI:** 10.1101/736843

**Authors:** Diana J. Rennison, Seth M. Rudman, Dolph Schluter

## Abstract

The processes of local adaptation and ecological speciation are often strongly shaped by biotic interactions such as competition and predation. One of the strongest lines of evidence that biotic interactions drive evolution comes from repeated divergence of lineages in association with repeated changes in the community of interacting species. Yet, relatively little is known about the repeatability of changes in gut microbial communities and their role in adaptation and divergence of host populations in nature. Here we utilize three cases of rapid, parallel adaptation and speciation in freshwater threespine stickleback to test for parallel changes in associated gut microbiomes. We find that features of the gut microbial communities have shifted repeatedly in the same direction in association with parallel divergence and speciation of stickleback hosts. These results suggest that changes to gut microbiomes can occur rapidly and predictably in conjunction with host evolution, and that host-microbe interactions might play an important role in host adaptation and diversification.

## Background

Bacteria play a crucial role in the physiology, ecology, and evolution of animals [1–5]. Like other interspecific interactions, the composition of affiliated microbial communities can impact host performance and relative fitness [6–9]. There is increasing appreciation for the role of gut microbiomes in the evolution of hosts [5, 10]. Recent studies have demonstrated that patterns of gut microbial community composition largely reflect host phylogeny (‘phylosymbiosis’ [11, 12]), that gut microbiomes can contribute to reproductive isolation [13, 14], and they can drive rapid evolution in host populations [15]. The importance of microbiomes on host performance has led to the suggestion that host evolution cannot be understood without consideration of their associated microorganisms [16, 17]. Despite the recognition of the importance of gut microbiomes in driving host evolution, relatively little is known about whether and how microbial changes affect local adaptation in natural host populations. This lack of knowledge stems in part from the inherent difficulty of studying the causes of adaptation and speciation in nature.

Cases of parallel evolution that are associated with repeated transitions in communities of interacting species have been vital in identifying how biotic interactions drive phenotypic and genomic change in nature [18–25]. Instances of parallel evolution might likewise be a useful tool for uncovering the relationship between host local adaptation and transitions in characteristics of gut microbial communities. If host evolution and gut microbiomes are linked, then a straightforward prediction is that local adaptation in hosts should be associated with repeatable changes in gut microbial communities [12]. These changes should be particularly repeatable in functionally important components of the gut microbiome, as microbial communities can exhibit parallelism in functional composition but not taxonomic composition [26]. Diet is a major factor that shapes the vertebrate microbiome [27–30], so cases of parallel evolution that led to convergence in diet create a particularly strong prediction of parallelism in the gut microbiome. Some prior work has assessed gut microbial divergence between ecotypes [31, 32], but little is known about whether parallel host evolution is associated with parallel changes in gut microbial communities.

Determining the strength of the association between parallel evolution in hosts and parallel shifts in gut microbial communities is a crucial step towards understanding the role of host-microbe interactions in the adaptive process. Sister taxa occurring sympatrically are ideal for understanding whether and how genetically-based differences in morphology, physiology, or behavior shape gut microbiomes in a natural setting, because the species inhabit a common environment. A common environment allows for an assessment of potential genetic differences in gut microbial communities in a natural context, where, unlike in most lab-rearing scenarios, heritable differences in host habitat choice and diet that influence gut microbial composition are expressed. If interactions with gut microbial communities have influenced the direction of host evolution or host evolution shapes host microbiomes, then we expect that features of microbial communities will reflect evolved differences between host populations in genotype and phenotype.

Threespine stickleback (*Gasterosteus aculeatus*) of British Columbia, Canada are an ideal system in which to uncover the relationship between host local adaptation and gut microbial communities in nature. Stickleback are a textbook case of repeated local adaptation. Marine stickleback independently colonized and adapted to many freshwater environments at the end of the last ice age ~11,000 years ago [33]. The adaptation of marine populations to freshwater conditions is characterized by parallel genetic, morphological, and physiological changes [33–38]. In five independent lakes, double colonization and natural selection has also driven evolution of sympatric pairs of stickleback species [39–41]. The two species within each pair differ greatly in their morphology and diet, and mate assortatively in the wild [19, 42, 43]. One is deep-bodied and forages in the nearshore environment primarily on aquatic larval insects and other invertebrates (termed the ‘benthic ecotype’). The other sympatric ecotype is shallow-bodied and forages primarily in the pelagic zone on zooplankton (termed the ‘limnetic ecotype’) [19, 43]. The repeated local adaptation and speciation in independent freshwater environments allows us to test the associations between environment, host ecotype, and gut microbial communities in nature.

To test for repeated changes in gut microbial community composition in association with parallel evolution we leverage the repeated evolution of benthic and limnetic stickleback ecotypes. We use both taxonomy and microbial functional to test whether microbiomes show parallelism across independently evolved species pairs. In an additional contrast we use the recent breakdown of one species pair into a population of advanced generation hybrids that are intermediate in morphology and diet (“reverse speciation”) [44–46] to examine whether the gut microbial community showed predictable changes. Finally, we use the marine to freshwater transition that many stickleback populations, including the sympatric species pairs, have undergone [33] to again test whether evolutionary parallelism leads to parallel gut microbial changes and to determine whether the diversity of gut microbial pathogens is reduced following colonization of a new habitat (*i.e.* the ‘honeymoon hypothesis’ [47]).

## Methods

### Field Collections

Field sampling of threespine stickleback from eight populations was done between April and May 2011, in the southwestern region of British Columbia, Canada (Supplementary Table 1). Five marine individuals were sampled from Oyster Lagoon, representing the ancestral marine population that founded these freshwater populations ~11,000 years ago [23, 33]. In three lakes containing species pairs (Little Quarry, Paxton and Priest) we sampled five individuals of each ecotype (benthic and limnetic; thirty individuals total). In addition, we sampled five individuals from Enos Lake, which contained a species pair of stickleback until they underwent reverse speciation in 2000 [45]. Fish were captured using unbaited minnow traps, which were set for 10-15 hours in high-usage foraging areas of each aquatic environment, prior to being euthanized with buffered MS-222 (Sigma, St Louis, MO, USA). To control for the effects of sex [48] and age, only adult male stickleback were included in the study.

### Lab Rearing

Three benthic x benthic, three limnetic x limnetic, and three benthic x limnetic crosses from Paxton Lake were raised in the common environment of the laboratory. Crosses were made in April 2011 using pure wild-caught parental fish. The offspring of these crosses were reared in 100 L fresh water tanks for 8 months; during this time, all crosses were fed the same diet of brine shrimp *Artemia, Mysis* shrimp, and chironomid larvae. At 8 months of age one male individual per family (9 individuals total) was euthanized using buffered MS-222 and prepared for DNA extraction.

### DNA extraction and sequencing

After euthanasia, the whole digestive tract was removed using sterile instruments. Any visible prey items within the digestive tract were removed. The posterior portion of the esophagus, the entire stomach, and the foremost anterior part of the intestine were triple rinsed using lysis buffer (~2ml/rinse) and all three rinses were combined. This lysis buffer solution was then immediately used in a standard phenol-chloroform DNA extraction protocol. The resulting DNA was amplified using the earth microbiome 515F-806R 16S rRNA primers [49], with three separate PCR amplification reactions per sample. The 3 PCR reactions from each sample were pooled, then uniquely barcoded and sequenced paired end (151bp x 151bp reads) on a MiSeq platform at the IGSB-NGS Core Facility at the Argonne National Laboratory in Lemont, Illinois USA. Negative controls were carried through the entire extraction process and showed no amplification.

### Bioinformatic analysis

MacQIIME version 1.9.1 was used for the analysis of raw illumina sequence reads to identify operational taxonomic units (OTUs), determine a phylogenetic tree, and calculate diversity metrics [50]. We followed the analysis pipeline outlined in Caporaso *et al*., [50], with the exception of the *denoiser* step, which was omitted as it is not required for Illumina data. Sequence reads were excluded if there was more than a one base pair error in the barcode. The pipeline began with trimming and demultiplexing of paired reads. Default cutoff values were used for quality score and read length. Chimeric sequences were then identified using ChimeraSlayer [51] and removed before performing downstream analyses. OTUs were picked using a 0.97 similarity threshold and the default parameters of the *pick_open_reference_otus* pipeline from MacQIIME (an open reference method). Reads identified as belonging to chloroplasts were removed from the resulting OTU table using the *filter_taxa_from_otu_table.py* script (and -n c__chloroplast parameter). The relative abundance of each taxa was estimated from the final OTU table, summarized at the level of order and plotted.

From the OTU table we calculated the core microbiome, which is a set of bacterial taxa shared by 100% (strict) or 60% (relaxed) of individuals using the *compute_core_microbiome.py* script in MacQIIME. The 100% and 60% core classifications indicate that the OTU is present in 100% or 60% of the individuals in a group respectively. We estimated the core microbiome for each ecotype (limnetic, benthic, hybrid, freshwater and marine). We then looked at the overlap of the core microbiome between each different ecotype.

To assess community composition and diversity, we rarefied the data 10 times to 90,000 sequences per sample (which resulted in the exclusion of two samples), used each independent rarefaction to estimate alpha (*alpha_diversity.py* script) and beta diversity (*jackknifed_beta_diversity.py* script), and then averaged the estimates across the 10 replicates (See Supplementary Figure 1 for rarefaction plots and Supplementary Table 2 for individual sequencing read counts). We estimated three alpha diversity metrics, richness, chao1 diversity, and phylogenetic diversity, for each sample. For beta diversity we estimated both Bray Curtis dissimilarity and unweighted UniFrac metrics [52], but as they produced largely concordant results we opted to focus on analyses using Bray Curtis dissimilarity. Singleton OTUs (those observed only once across the dataset) were removed from the rarefied dataset before further downstream analysis. The final OTU dataset consisted of 14,991 OTUs.

Microbial function assignments were done using PICRUSt version 1.1.0 [53]. Since the OTUs for the above analyses were picked using an open reference it was necessary to filter the OTU table so that it only contained OTUs found in the Green Genes database [54]. Filtering of the OTU table was done against Green Genes 13.5 release with 97% sequence similarity threshold. The filtered OTU table was normalized by copy number, then used to predict the meta-genome with estimation of NSTI scores. The mean and median NSTI score for our samples was 0.08 (range 0.06 - 0.11 across populations). We were then able to categorize the OTUs by biological function, yielding KEGG annotations.

### Statistical analysis

To determine whether stickleback ecotypes were associated with differences in microbiome alpha diversity we used linear mixed effects models to examine species richness with fish ecotype as a fixed effect and lake of origin as a random effect. Gut microbial community composition was quantified using Bray-Curtis dissimilarity between individuals based on estimated microbial abundance. Dissimilarity values were analyzed with NMDS in R using the *Vegan* package [55]. Only the first 5 NMDS axes were retained in subsequent tests of parallelism and when examining the effects of freshwater colonization. Separate NMDS analyses were carried out to test for parallelism in taxonomic composition (OTUs) and microbial community function (KEGG designation). Independent evolutionary events of speciation and freshwater colonization were used as statistical replicates for all tests. Results were visualized using the *ggplot2* package [56].

To quantify parallelism in gut microbial composition and function we measured the angles between multivariate vectors based on the first 5 NMDS axes that describe the direction of the difference between populations in their gut microbiomes [57]. Each vector represents the direction and magnitude of divergence in gut microbiome composition (taxonomic diversity or function) between ecotypes (benthic and limnetic or marine and fresh). A small angle between the divergence vectors of two ecotype pairs represents a high degree of parallelism in gut microbial divergence. A 90° angle would indicate no parallelism in the pattern of gut microbial divergence, and a large angle (closer to 180°) indicates a dissimilar direction of divergence. This vector based approach has previously been used to estimate parallelism in phenotypes and genotypes between populations diverging repeatedly across similar environments [e.g., 58]. We described the direction of divergence between ecotypes within each pair using a vector connecting the mean position (centroid) of individuals of one ecotype (e.g., limnetic) to the mean position of individuals of the other ecotype (e.g., benthic). We estimated the angle (θ, in degrees) between divergence vectors of each ecotype pair (from each lake) and calculated the average angle. To assess parallelism, we then tested whether the average angle between divergence vectors of different ecotype pairs was smaller than expected by chance. We used t-tests to determine significance and 90° as the null or random expectation.

To understand the taxonomic and functional underpinnings of any observed (or lack of) parallelism we assessed differences between host populations in the abundance of specific taxonomic and functional microbial groups. We compared the relative abundance of each of the 87 bacterial orders between ecotypes in each population comparison (benthic vs. limnetic fish, marine vs. freshwater fish, and enos hybrids vs. either pure ecotype). We highlight the taxonomic groups where we found the largest differences in abundance between the stickleback ecotypes (i.e. those falling in the 10th and 90th quantiles for difference in abundance. We tested for significant differences between ecotypes (Benthic-Limnetic, Marine-Fresh. Hybrid-Benthic, Hybrid-Limnetic) using Kruskal-Wallis tests for the relative abundance of microbes falling into the 41 KEGG gene function categories. Relative abundance for gene function is defined as the percent of the predicted metagenome made up of a given KEGG functional module (category). The P-values resulting from each set of tests were corrected for multiple testing using the BH method [59] with the *p.adjust* function.

We also assessed the extent of gut microbial divergence between ecotypes measured as distance in NMDS coordinates. We extracted NMDS coordinates for each individual within each population. We then calculated the average pairwise Euclidean distance between individuals within a population and compared this distance to the average pairwise distance between individuals in different populations. For the benthic-limnetic comparisons we determined the average pairwise distance within and between ecotypes for each of the three independent species pairs separately and then averaged them. To assess how reverse speciation in Enos Lake altered the gut microbial community, we calculated the average of pairwise distances between individuals from Enos Lake and contrasted that with distance between these Enos Lake individuals and all individuals from each benthic and limnetic population. For the marine-freshwater contrast we compared variation between individuals within each freshwater population to variation between individuals from each freshwater population and individuals from the marine environment. To assess whether individuals from different ecotypes showed significant differences in gut microbiome composition, we conducted MANOVAs on the first 5 NMDS axes (to match parallelism analysis) with ecotype as a fixed effect.

To test whether more recently colonized freshwater populations carried lower pathogenic loads (i.e. the ‘honeymoon hypothesis’), we estimated and compared the relative abundance of the bacterial families known to be pathogenic in fish between marine and freshwater ecotypes. Bacterial taxa were classified as pathogenic if they were identified as belonging to a bacterial family previously shown to cause disease in wild or farmed fish (Austin and Austin, 2016). A two-sample t-test was used to test for a statistical difference in average pathogenic load between marine and freshwater fish.

## Results

### Parallel shifts in function but not composition of microbial communities with repeated ecological speciation

We found considerable parallelism in the direction of the difference between independent limnetic and benthic ecotype pairs in the functional properties of the gut microbial communities. The average angle was 17.45° between divergence vectors for the three species pairs, which is significantly more parallel than expected by chance (*P* = 0.012, *T_2_*=-23.71). In contrast, the average angle between divergence vectors based on taxonomic composition shows substantially less parallelism and was not quite significantly different from the random expectation (76.4°, *P* = 0.06, *T_2_*=-4.303) (Figure 1A).

**Figure 1.**
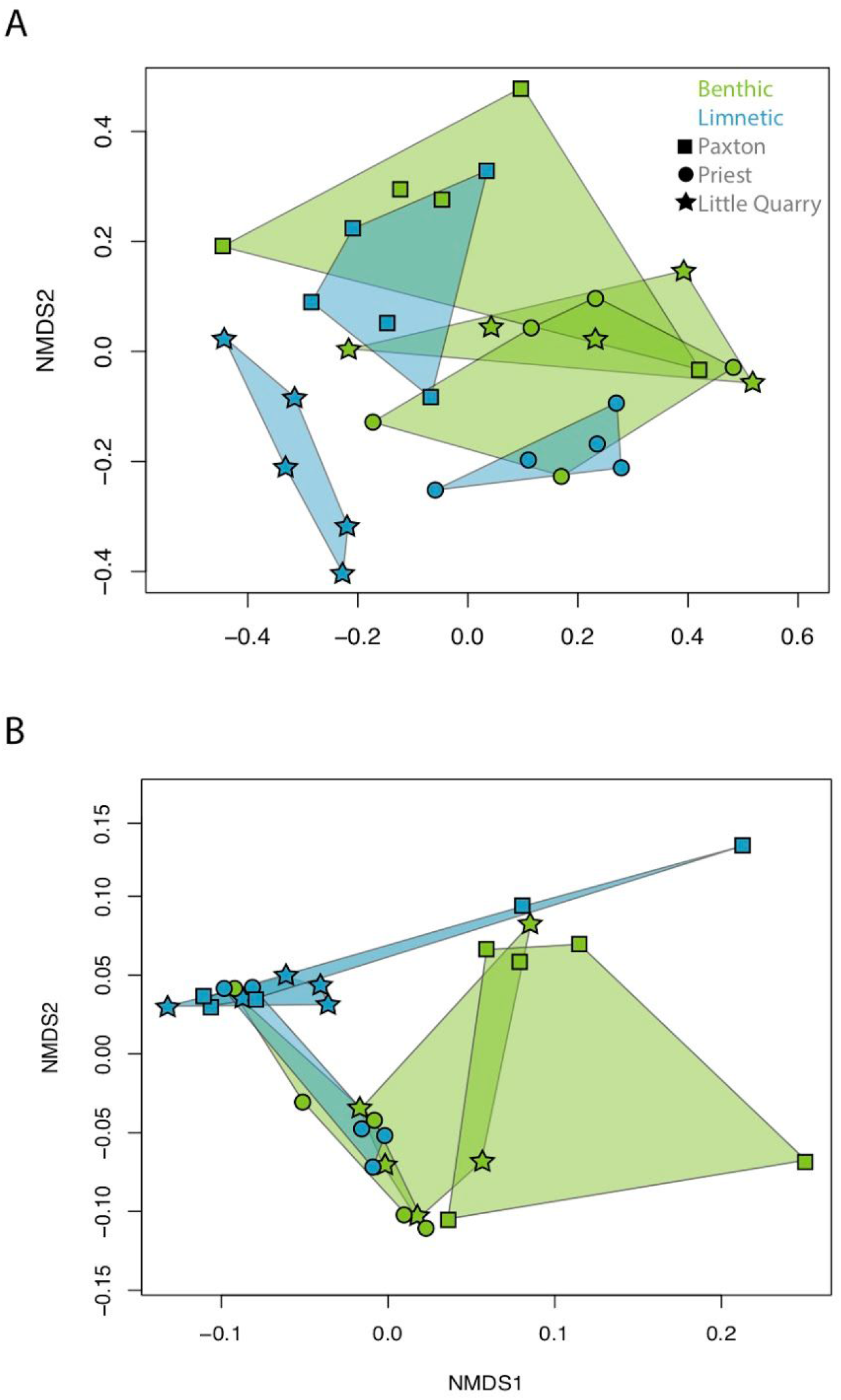
Differentiation of the (a) taxonomic composition and (b) functional composition of the gut microbiome of benthic and limnetic threespine stickleback from Paxton, Priest and Little Quarry based on Bray-Curtis dissimilarity.

Comparison of the proportion of the metagenome made up of a given KEGG functional module suggested several consistent differences between the microbiomes of benthic and limnetic ecotypes in multiple functional categories (Figure 2A). After correction for multiple testing fifteen KEGG functional modules differed significantly (p < 0.05) in relative abundance between benthic and limnetic ecotypes (indicated with asterisks in Figure 2A); eight of the differentially enriched categories pertained to metabolism.

**Figure 2.**
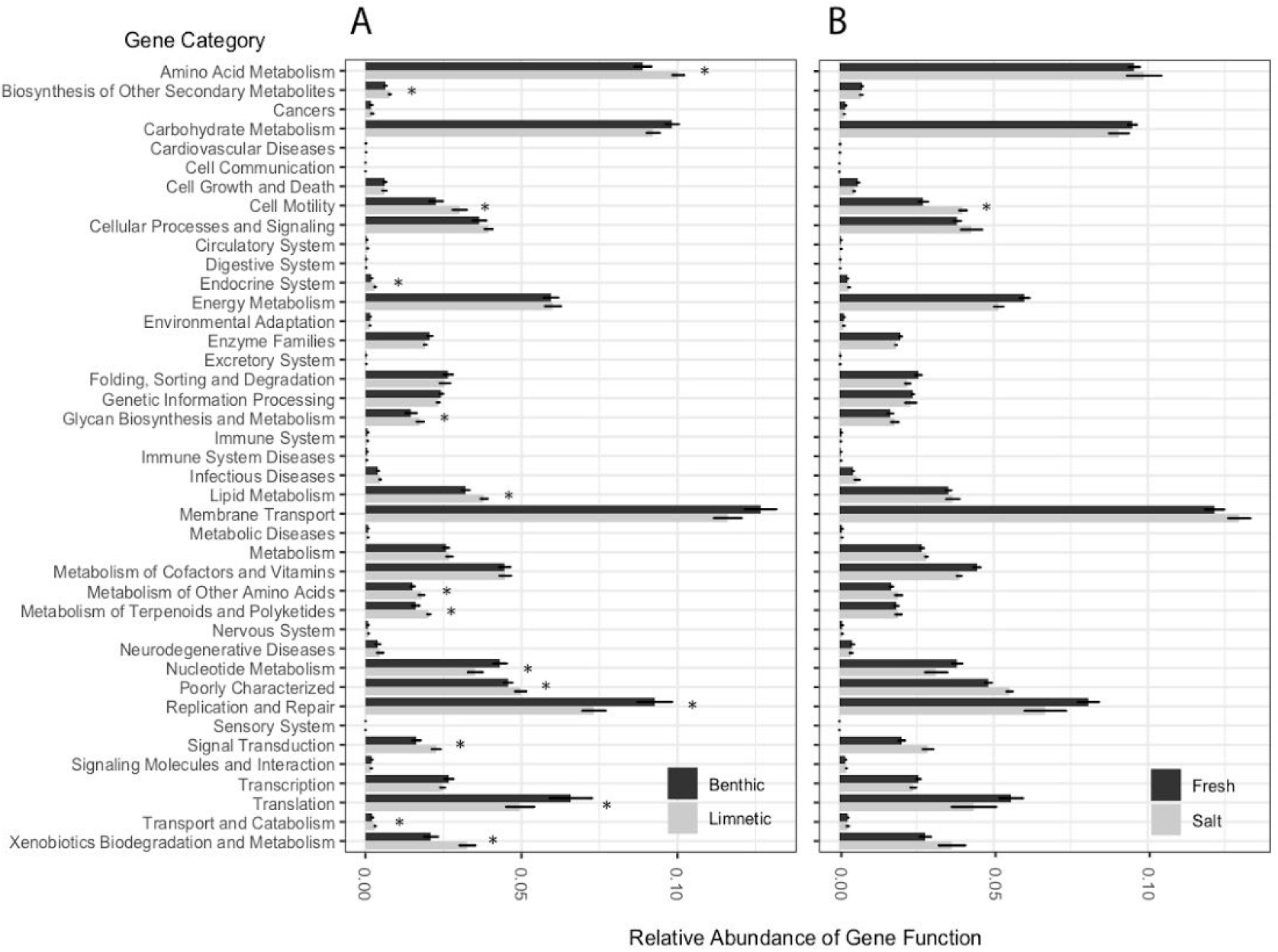
Results of PICRUSt analysis showing relative abundance of KEGG orthologs for stickleback (a) benthic and limnetic ecotypes and (b) marine and fresh ecotypes. Significant differences in abundance are indicated with an asterix (P < 0.05 after correction for multiple testing).

Although we did not observe parallelism in the vectors based on taxonomic composition there were orders that were notably more enriched in each ecotype. Six orders (Acidimicrobiales, Caulobacterales, Chroococcales Gloeobacterales, Rhodospirillales and Pseudanabaenales) were on average 20-492x more abundant in limnetics than benthics and bacteria belonging to Neisseriales were on average 32x more abundant in benthic than limnetic fish. Using the relaxed criterion for the composition of the core microbiome, we identified 212 core OTUs in the core microbiome of wild benthics (3.3% of the 6347 OTUs found across all the benthic samples) and 282 core OTUs for wild limnetics (4.1% of the 6923 OTUs found across all limnetic samples). 49% of these core OTUs were shared between the benthic and limnetic ecotypes cores. 168 OTUs were unique to either benthics or limnetics; the majority of unique OTUs were proteobacteria or firmicutes, belonging primarily to one of four bacterial orders (Burkholderiales (20%), Clostridiales (14%), Pseudomonadales (19%) or Sphingomonadales (11%)). Using the stringent criterion, wild benthic fish had 43 OTUs in their core microbiome and limnetics had 60 OTUs. There was a 45% overlap of the OTUs from the stringent core microbiome of limnetics and benthics, with the overlapping taxa largely coming from Gammaproteobacteria (13 of the 32 OTUs). Using the stringent criterion there were 39 OTUs that were unique to either ecotype. Again, these primarily belonged to Pseudomonadales (42%), Burkholderiales (15%) and Sphingomonadales (18%)(See Figure 3 for relative abundance of bacterial orders for each ecotype).

**Figure 3.**
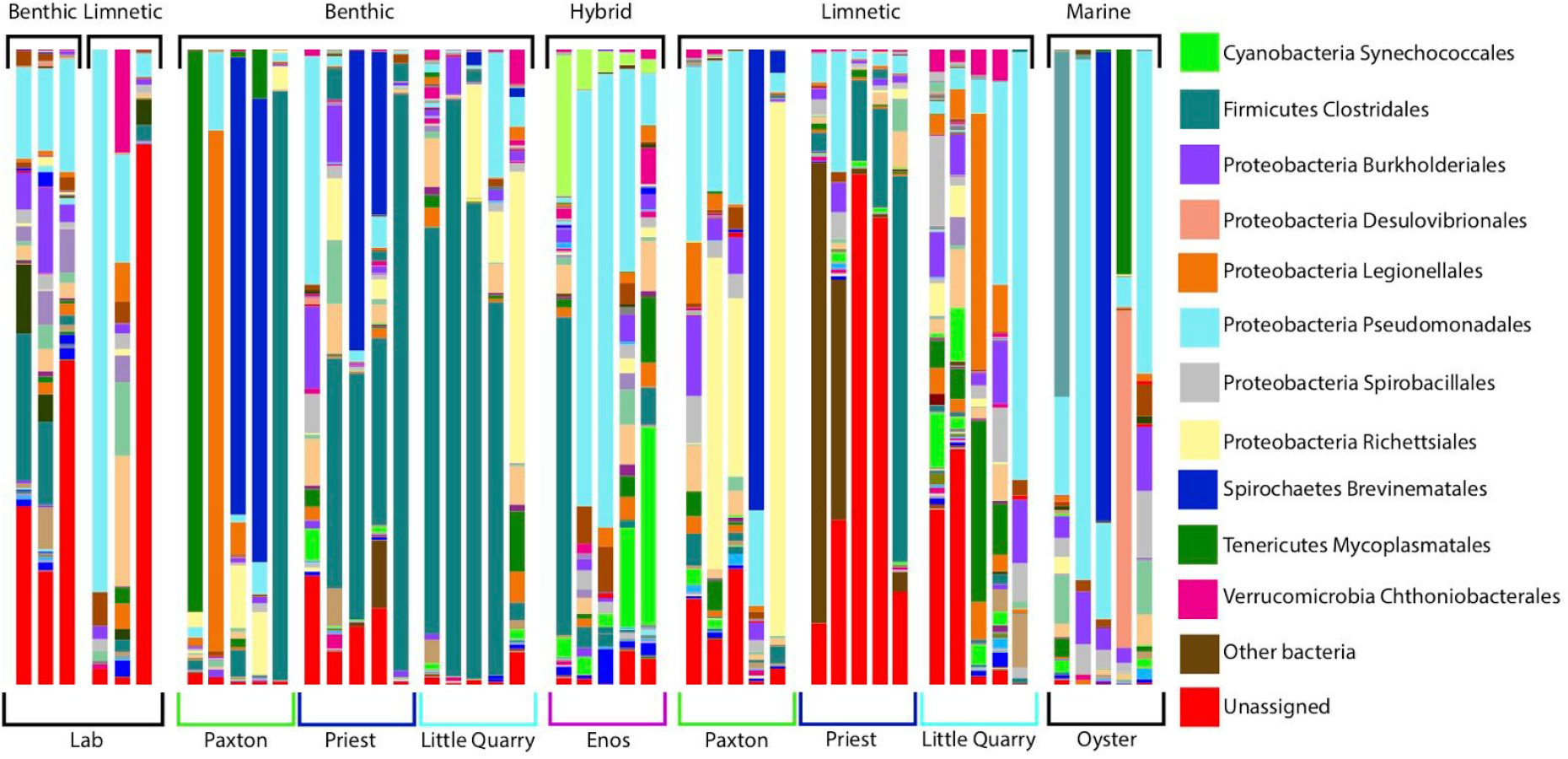
Taxonomic composition of the gut bacterial communities of stickleback individuals as grouped by ecotype and lake of origin. Abundant microbial groups are classified to order or class by color. Bar height indicates fraction of relative abundance.

The composition of the gut microbial community was more similar between individuals from the same ecotype (distance: 0.33, sd=0.10) than between individuals from different ecotypes (distance: 0.42, sd=0.15), based on distances in ordinal (NMDS) space when we compared across all lakes (Figure 1). This similarity in gut microbial composition between individuals of the same ecotype was more pronounced in the limnetic ecotype and less so in the benthic (limnetic: x=0.24, sd=0.03, benthic: x=0.42, sd=0.06) which was driven by reduced variance within lakes in limnetic populations relative to benthic populations. Across individuals from all pairs ecological speciation was associated with a significant shift in the gut microbiome taxonomic composition of benthic and limnetic stickleback ecotypes (F_1,22_=3.08, p=0.030, Table S3). There was no significant difference between benthic and limnetic ecotypes in gut microbial species richness (F_1,25_=0.54, p=0.54). When reared in the lab environment we did not observe any evidence that individuals from the same ecotype had more similar gut microbial communities than individuals from different ecotypes (within: 0.55 (sd=0.21), between: 0.524 (sd=0.13)) (Supplementary Figure 2).

### When reproductive isolation breaks down, the gut microbiome also changes

The breakdown of reproductive isolation in Enos Lake (reverse speciation) largely led to a microbiome community that was intermediate in both composition and function relative to extant benthic and limnetic individuals (Figure 4). The gut microbiome composition of Enos hybrid fish was significantly different in composition than that found in extant benthic fish (F_1,12_=4.22, p=0.019, Table S3) and showed a trend towards divergence from the community found in extant limnetic fish (F_1,12_=3.06, p=0.052, Table S3). Gut microbe community composition differed less among Enos Lake hybrids than when compared to either benthic or limnetic individuals from intact species pairs (within: 0.228 (sd=0.18), benthic: 0.416 (sd=0.21), limnetic=0.316 (sd=0.16)). The aspects of the gut microbiome community of Enos Lake individuals that were different or unique relative to the intact species pairs includes ten orders (Actinomycetales, Bdellovibrionales, Chroococcales, Chromatiales, Gaiellales, Methylococcales, Neisseriales, Pseudanabaenales, Solirubrobacterales and Synechococcales) that were enriched 23-586x in the hybrid fish relative to intact benthics. Three orders were enriched 50-429x in the Enos hybrids relative to intact limnetics (Caldilineales, Chromatiales and Neisseriales). There were also two bacterial orders present in the Enos hybrids (Acidobacteriales and Methanocellales) that were completely absent from any of the intact limnetics or the intact benthics.

**Figure 4.**
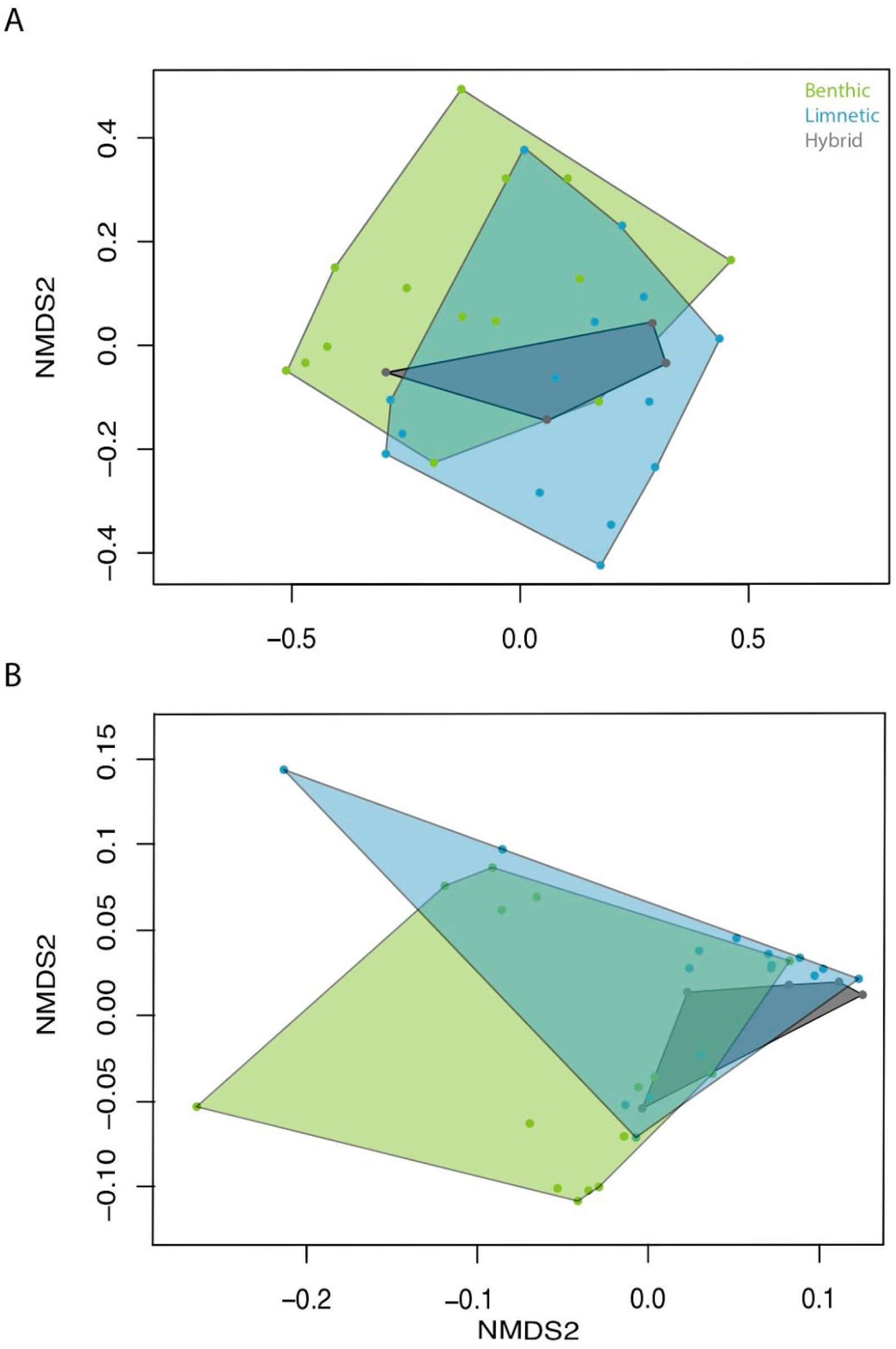
Differentiation of the (a) taxonomic composition and (b) functional composition of the gut microbiome of hybrid threespine stickleback from Enos lake relative to benthic and limnetic stickleback from Paxton, Priest and Little Quarry based on Bray-Curtis dissimilarity.

Enos hybrids had microbiomes that were largely intermediate in function relative to intact benthics and limnetics (Supplementary Figure 3). After correction for multiple testing there were no significant differences (P > 0.05) in the abundance of taxa associated with any of the KEGG gene categories when comparing Enos fish to intact limnetics. Between Enos fish and benthics there were trends (0.1 > P > 0.05 after multiple testing correction) towards differences in functional abundance, with nine categories enriched in Enos hybrids and five enriched in benthics (indicated by asterisks in Supplementary Figure 3). Most of the relevant functional categories of these trends related to metabolism or biosynthesis.

A comparison of the core microbiome of Enos hybrids with those of the extant species pairs also revealed unique aspects of the hybrid microbiome. There were 345 OTUs for Enos using the relaxed criteria (9.8% of the 3512 total OTUs found in Enos), and 115 with the strict criteria. The core microbiome of the Enos hybrids overlapped to the same degree with the pure benthic fish from the other three lakes (41% overlap of OTUs using the relaxed criteria; 39% for strict criteria) as with the pure limnetics (41% overlap of OTUs using the relaxed criteria; 38% for strict criteria). Using the relaxed criteria, 30% of OTUs were unique to the Enos hybrids (45% using the strict criteria). Most unique taxa were Cyanobacteria (15%), Planctomycetes (19%) or Proteobacteria (49%) There was no significant difference between hybrids and extant species pairs in gut microbial species richness (F_2,4.65_=0.57, p=0.59, Figure S4).

### Freshwater colonization-differentiation and a test of the honeymoon hypothesis

The direction of divergence in taxonomic composition of the gut microbiome from the marine ancestral population to each of the seven freshwater populations was significantly parallel. On average, the angle (θ) between the divergence vectors of pairs of marine and freshwater populations was 38.18°, which is less than half the null expectation of 90° (*P* < 0.0001, *T_5_*=-14.711) (Figure 5A). Individual fish from marine and freshwater populations differed significantly in their gut microbiome composition (F_1,31_=4.28, p=0.004, Table S3). As predicted by the ‘honeymoon hypothesis’, freshwater populations, on average, had a significant reduction in the abundance of pathogenic taxa relative to the marine population (*P*=0.003, *T_6_*= 4.71), with an average of 16% fewer bacteria belonging to pathogenic genera. Divergence in the functional composition of the gut microbiome across the freshwater populations was more weakly parallel, with an average angle of 49.69° (*P* = 0.003, *T_5_*=-5.41).

**Figure 5.**
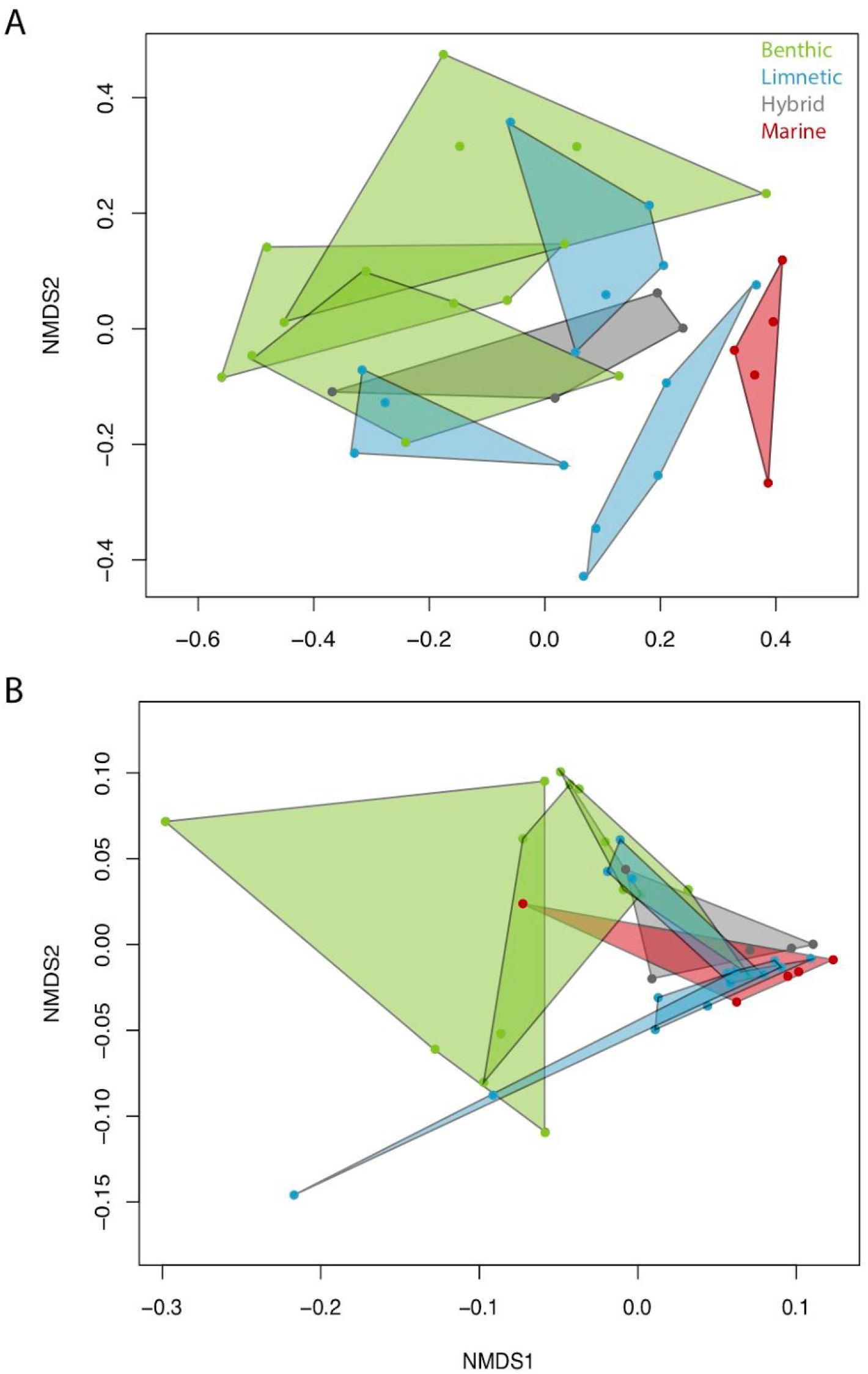
Differentiation of the (a) taxonomic composition and (b) functional composition of the gut microbiome of marine and freshwater threespine stickleback based on Bray-Curtis dissimilarity.

The parallel divergence in gut microbial community of populations found in freshwater was explained, in part, by large differences in the relative abundance of several orders. Of the orders found in both population types, three orders were substantially enriched (29 - 334x) in marine fish relative to freshwater fish (Desulfovibrionales, Mycoplasma and Vibrionales). Eight orders had a 20-665x enrichment in freshwater fish (Chthoniobacterales, Clostridiales, Cytophagales, Flavobacteriales, Gemmatales, Planctomycetales, Rickettsiales, and Xanthomonadales). Similarly, analysis of the proportion of the metagenome made up of a given KEGG functional category revealed differences between marine and freshwater stickleback in the abundance of several categories, although only cell motility was significantly different after correction for multiple testing. The categories most considerably enriched in marine stickleback relative to freshwater populations were those associated with amino acid metabolism, cell motility, membrane transport, signal transduction, and xenobiotics biodegradation & metabolism. Among freshwater fish, the categories most considerably enriched were functions associated with carbohydrate metabolism, energy metabolism, replication and repair, and the metabolism of cofactors and vitamins (Figure 5B).

The core microbiome of marine stickleback had 453 OTUs using the relaxed criteria (22.5% of the 2014 OTUs identified in the Marine samples) and 132 OTUs using the stringent criteria. The core microbiome of freshwater fish had 222 OTUs using the relaxed criteria and 31 OTUs using the stringent criteria. Using the relaxed criteria, the core microbiome of marine and freshwater fish had a 35% overlap (20% for the strict criteria); nearly half of the overlapping OTUs using both the strict and relaxed criteria were Gammaproteobacterial orders. The taxa unique to the marine fish were largely Psuedomonadales (61%). Microbiomes of marine and freshwater stickleback were distinct, as we found that variation in gut microbial communities between marine and freshwater individuals exceeded the variation found between freshwater individuals (within freshwater: 0.31 (sd=0.07), between freshwater and marines: 0.49 (sd=0.15)). There was no significant difference between marine and freshwater stickleback populations in gut microbial species richness (F_1,3.88_=1.05, p=0.36, Figure S4).

## Discussion

### Parallel host evolution and parallel shifts in gut microbial communities

We found evidence that independently evolved benthic and limnetic stickleback ecotype pairs show parallel changes in their gut microbiomes. This study is the first to explicitly test for parallel changes in gut microbial composition or function using cases of repeated diversification. Prior studies examining the relationship between parallel host evolution and host gut microbiomes grouped independently evolved populations and used individuals as replicates in their tests of parallelism [31, 32]. While previous studies demonstrated that local adaptation can alter gut and kidney microbial communities, they found little evidence that differences in microbiomes are parallel across independent cases of host adaptation [31, 60]. For example, Trinidadian guppies (*Poecilia reticulata*), a well-studied system for parallel phenotypic evolution [61–63], have different microbial community composition in upstream and downstream populations but these patterns appear not to have evolved in parallel across watersheds [31].

Host diet and host genotype are the most likely causes of the parallel shifts we observed in microbiome composition across stickleback species pairs. Independent benthic and limnetic ecotypes exhibit parallel shifts in diet and there are substantial differences in diet between ecotypes [19, 43]. Diet has previously been shown to strongly influence gut microbial communities in a variety of species [12, 64–66], including stickleback [29]. Benthic and limnetic individuals raised in a lab common garden and fed a common diet did not show substantial differences in microbial composition, reinforcing that diet may be an important factor in microbiome differentiation. Independently evolved ecotypes also show parallel genomic evolution, including remarkable parallelism in SNPs linked with genes that influence immune function [23]. The variation in diet between stickleback individuals also has a strong genetic basis [67]. As such, parallel allelic changes across replicate populations could contribute to the parallel differences we observed in gut microbial communities, as host genotypes strongly impact microbiome community composition [9, 68]. Future work that uses reciprocal transplants within a natural environment, constraining each ecotype in both the nearshore or open water environment to constrain their diet, could help to disentangle the relationship between host diet and host genotype in driving gut microbial composition. Large-scale individual-level sequencing of microbiomes and host genotypes from genetically diverse host populations also presents the opportunity to find correlations between specific SNPs, windows, or haplotypes and variation in gut microbiome composition.

Stickleback species pairs provide a potentially useful system for further investigation of how host-microbe interactions shape organismal performance, local adaptation, and speciation. Parallelism across pairs of ecotypes was more pronounced in microbial function than taxonomic composition, suggesting that divergence between ecotypes may influence organismal performance and, ultimately, fitness. The gene functional categories of metabolism and biosynthesis were most differentiated between ecotypes, and previous work has found substantial population-level variation in stickleback metabolism [69, 70]. KEGG functional inferences are based on the similarity of OTUs in a sample to those with known functions. 2-3% sequence divergence across microbial OTUs is typically considered to reflect species level differences. Our KEGG functional inferences of OTUs were based on comparisons with an average of 8% divergence from reference taxa with known functions (based on NSTI scores). As such, it is possible that we are over- or under-estimating the degree of parallelism when using predicted KEGG functions. Measuring the contribution of the functional differences between ecotypes to fitness and ongoing ecological speciation presents some challenges. Assessing whether variation in the gut microbial community and function is correlated with differences in fitness within each ecotype could provide some correlative information on the links between gut microbial composition and fitness. If the gut microbial differences we observed between ecotypes were found to enhance the fitness of each ecotype in its preferred environment, but not in the alternative environment, the gut microbiome could be implicated in maintaining reproductive isolation. Gut microbial communities can influence mate choice decisions [13], and the differences we observed between ecotypes could be a component of reproductive isolation by contributing directly to mate choice. Lab-based mate choice trials are useful in assessing reproductive isolation in stickleback [71] and manipulating the gut microbial community to test for effects of mate choice is feasible. These types of approaches assessing the relationship between microbiome composition and host evolution will be important for understanding the role that microbiomes play in the genesis of host diversity [10, 13, 15, 17].

### Speciation, reverse speciation, and shifts in the gut microbial community

Phenotypic evolution of keystone or dominant species can alter ecological patterns and processes [72], including community composition [73, 74]. Ecological speciation in stickleback has been shown to impact community structure and ecosystem functions, with sympatric ecotypes having particularly strong effects on the prey community [75–77]. Our data demonstrate that the ecological effects of speciation also extend to repeated shifts in gut microbial communities (Figure 1). Previous work on the ecological consequences of evolutionary change in stickleback suggests that the some of the consequences are predictable based on the direction of evolution of functional traits [46]. Here we find similar patterns, with gut microbiomes showing divergence between benthic and limnetic ecotypes and with hybrid fish from Enos lake being intermediate and somewhat distinct from what is found in the extant pure limnetic and benthic ecotypes. This mimics the morphological evolution associated with speciation and reverse speciation across populations of stickleback. The general concordance between changes in gut microbiome and evolution suggests that gut microbiomes may shift both in a predictable direction relative to other populations based on phenotypic evolution and diet shifts. Lab reared F1 crosses between ecotypes did not show the same pattern of intermediate microbiome composition (Supplementary Figure 4), suggesting diet shifts as a prominent component of concordance between evolution and microbiome compositional shifts in nature. Similar comparative approaches in other taxa would be useful in determining whether concordance between morphological evolution and shifts in gut microbiome community composition is a general finding or specific to elements of divergence in stickleback.

### Colonization of new environments and the gut microbiome

Marine stickleback have colonized many freshwater environments independently, yielding to a well-replicated natural experiment on adaptation to freshwater environments [33]. Adaptation to freshwater involves substantial phenotypic and genomic parallelism, with a set of loci and inversions associated with adaptation to freshwater across independent populations. Marine populations colonizing freshwater environments face several physiological challenges [37] and changes in the microbiome could be a component of both acclimation and adaptation. We surveyed seven freshwater populations and found that the combination of freshwater colonization and adaptation to the freshwater environment, has driven shifts in the gut microbiome. A previous meta-analysis across fish species documented that marine and freshwater fish often differ in their gut microbial communities [64]. Our results add to this finding, as we also found evidence that shifts in taxonomic composition were parallel among independently derived freshwater populations relative to their marine ancestors. Although previous work suggests that variation in environmental microbes is not a major factor influencing the microbiome of stickleback [30]. Our data do not allow us to examine the influence of differences in the microbial environment across aquatic habitats on stickleback microbiomes, as unfortunately we did not collect environmental microbial samples taken from each aquatic environment. Without a comprehensive survey of the microbial environments in marine and freshwater it is difficult to determine the relative contribution of differences in the microbial environment and host genotype to microbiome composition. Translocation experiments, particularly reciprocal transplants, where marine fish are reared in freshwater and freshwater fish are reared in marine environments, could be used to determine the relative contribution of environment and host adaptation on the observed parallel shifts in microbiome composition of freshwater stickleback populations.

A component of the microbial differences between marine and freshwater stickleback is reduced gut microbial pathogen loads, as all of the surveyed freshwater populations had lower relative abundances of putatively pathogenic taxa than marine populations. Our results are consistent with previous work demonstrating that pathogen load and diversity decreases after host colonization of a novel environment and pathogen load lag during range expansion [47]. The reduction of pathogens when colonizing this novel freshwater environment could provide an energetic benefit that enhances fitness and increases the probability that colonizing populations persist. Future work to assess how quickly pathogens are lost and the size of the fitness benefit for the host population could provide insight into the dynamics of adaptation to novel environments. More broadly, experimental work that tracks how gut microbial communities change as local adaptation occurs could prove useful for understanding the role that host-microbe interactions play in host adaptation.

### Conclusions and outlooks for the future

Our results uncover a link between host adaptation, speciation, and the gut microbiome. There is clear evidence in stickleback that resource competition has driven genetically determined shifts in diet and trophic morphology [43, 67, 78] and that this evolution can have effects on the ecology of the ecosystems in which stickleback occur [46, 75, 77, 79]. Our findings suggest that this phenotypic evolution is also associated with ecological changes in the gut microbiome of these fish. Moreover, we detect parallel changes in the microbiome across evolutionary independent host lineages evolving in parallel. This concordance between host evolution and microbiome divergence creates an opportunity for future work to test the factors that drive this pattern and to determine how host-microbe interactions shape local adaptation and speciation. Particularly enticing and tractable are field translocation experiments that could be used to tease apart the effect environment, diet, and host genotype in driving patterns of gut microbial divergence between populations. This work would help build towards a more comprehensive understanding of the frequency and magnitude with which host-microbe relationships influence host adaptation could be transformative to our understanding of the process of local adaptation.

## Ethics

Collections were made under the Species At Risk Act collection permit 236 and British Columbia Fish Collection permit NA-SU12-76311. Animals were treated in accordance with University of British Columbia Animal Care protocols (Animal Care permit A11-0402)

## Data accessibility

The resulting sequencing data archived in the NCBI SRA under Bioproject PRJNA475955. The R code has been archived in the Dryad database [doi:10.5061/dryad.m8p3q04].

## Author Contributions

DJR and SMR conceived of the project, carried out field collections, molecular work, analysis, and wrote the manuscript together. DS advised the project at all stages and helped write the manuscript.

