## Supplemental Materials for "Parallel changes in gut microbiome composition and function in parallel local adaptation and speciation"

**Supplementary Materials:**

Supplementary Table 1: Sampling Information

| Location | Population Types | Number of Individuals |
|---|---|---|
| Little Quarry, Nelson Island | Benthic and Limnetic | 5 Benthic<br>5 Limnetic |
| Paxton Lake, Texada Island | Benthic and Limnetic | 5 Benthic<br>5 Limnetic |
| Priest Lake,<br>Texada Island | Benthic and Limnetic | 5 Benthic<br>5 Limnetic |
| Enos Lake, Vancouver Island | Hybrid | 5 |
| Oyster Lagoon | Marine | 5 |

Supplementary Table 2: Sequencing Depth Information

| **Sample Name** | **Number of Reads** |
|---|---|
| **Enos_1** | 263785 |
| **Enos_2** | 162060 |
| **Enos_3** | 221898 |
| **Enos_4** | 52433 |
| **Enos_5** | 119908 |
| **F1_1** | 249198 |
| **F1_2** | 286382 |

| | |
|---|---|
| **F1_3** | 258084 |
| **Lab_Benthic_1** | 199326 |
| **Lab_Benthic_2** | 254145 |
| **Lab_Benthic_3** | 174560 |
| **Lab_Limnetic_1** | 229710 |
| **Lab_Limnetic_2** | 338744 |
| **Lab_Limnetic_3** | 135637 |
| **LQ_Benthic_1** | 386895 |
| **LQ_Benthic_2** | 193048 |
| **LQ_Benthic_3** | 317028 |
| **LQ_Benthic_4** | 354577 |
| **LQ_Benthic_5** | 8517 |
| **LQ_Limnetic_1** | 311149 |
| **LQ_Limnetic_2** | 279873 |
| **LQ_Limnetic_3** | 210474 |
| **LQ_Limnetic_4** | 282915 |
| **LQ_Limnetic_5** | 130182 |
| **Oyster_1** | 177604 |
| **Oyster_2** | 255892 |
| **Oyster_3** | 351335 |
| **Oyster_4** | 211424 |

| | |
|---|---|
| **Oyster_5** | 98692 |
| **Pax_Benthic_1** | 295771 |
| **Pax_Benthic_2** | 269054 |
| **Pax_Benthic_3** | 206654 |
| **Pax_Benthic_4** | 333619 |
| **Pax_Benthic_5** | 350804 |
| **Pax_Limnetic_1** | 206183 |
| **Pax_Limnetic_2** | 212249 |
| **Pax_Limnetic_3** | 175808 |
| **Pax_Limnetic_4** | 353157 |
| **Pax_Limnetic_5** | 193084 |
| **Pri_Benthic_1** | 293703 |
| **Pri_Benthic_2** | 253183 |
| **Pri_Benthic_3** | 283444 |
| **Pri_Benthic_4** | 136522 |
| **Pri_Benthic_5** | 293976 |
| **Pri_Limnetic_1** | 245312 |
| **Pri_Limnetic_2** | 279793 |
| **Pri_Limnetic_3** | 301799 |
| **Pri_Limnetic_4** | 286803 |
| **Pri_Limnetic_5** | 357676 |

Supplementary Table 3: Results from community composition analysis using both Bray-Curtis and weighted Unifrac diversity metrics. To match with analyses of parallelism all tests were conducted MANOVAs on the first 5 NMDS axes.

| contrast | ecotypes | metric | DF | F | p-value |
|---|---|---|---|---|---|
| Effect of ecological speciation | Benthic-Limnetic | Bray | 1,22 | 3.08 | 0.03 |
| | | Unifrac | 1,22 | 2.93 | 0.036 |
| | Benthic-Limnetic (Lab) | Bray | 1,5 | 0.5 | 0.65 |
| | | Unifrac | 1,5 | 0.68 | 0.57 |
| Community effects of reverse speciation | Benthic-Enos | Bray | 1,12 | 4.22 | 0.019 |
| | | Unifrac | 1,12 | 2.74 | 0.07 |
| | Limnetic-Enos | Bray | 1,12 | 3.06 | 0.052 |
| | | Unifrac | 1,12 | 9.61 | 0.0007 |
| Effect of freshwater colonization | Marine-Freshwater | Bray | 1,31 | 4.28 | 0.004 |
| | | Unifrac | 1,31 | 5.37 | 0.001 |

Supplementary Figure 1. Rarefaction plots for each individual indicating estimated alpha diversity for the number of sampled sequencing reads.

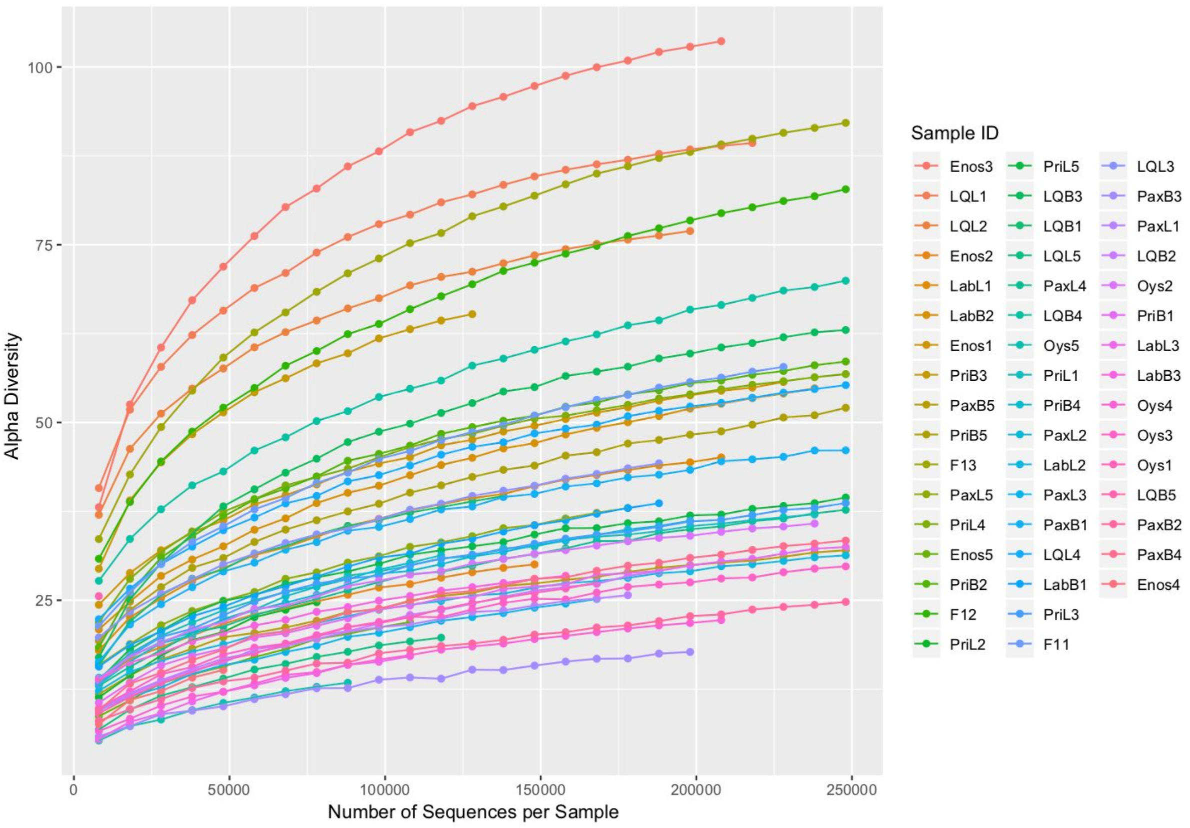

Supplementary Figure 2: Alpha diversity metrics for gut microbial communities from each stickleback ecotype. Black points show the means (+/-SEM) for each ecotype, colored points show the mean (+/-SEM) for each population within each ecotype. Panel A - species richness, Panel B - Chao1, Panel C - Phylogenetic diversity.

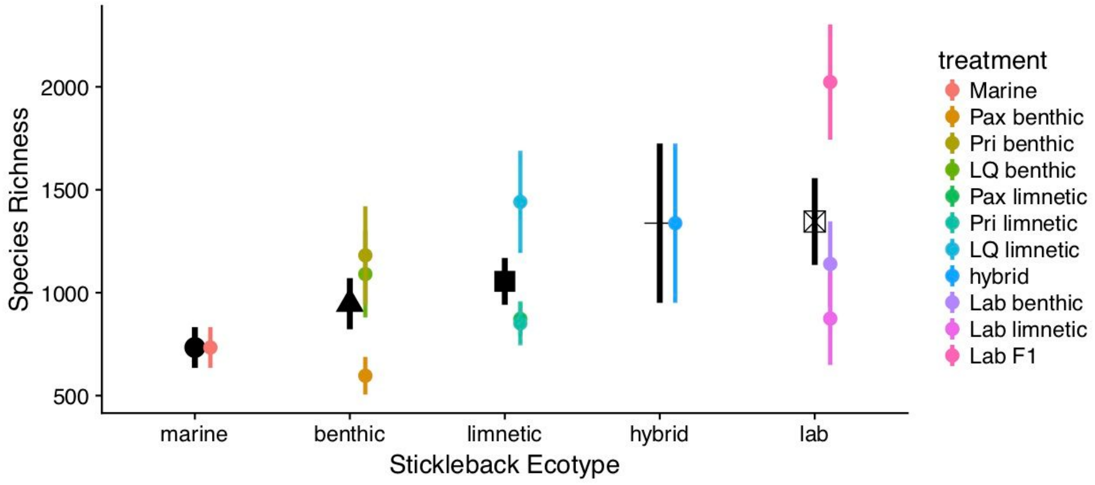


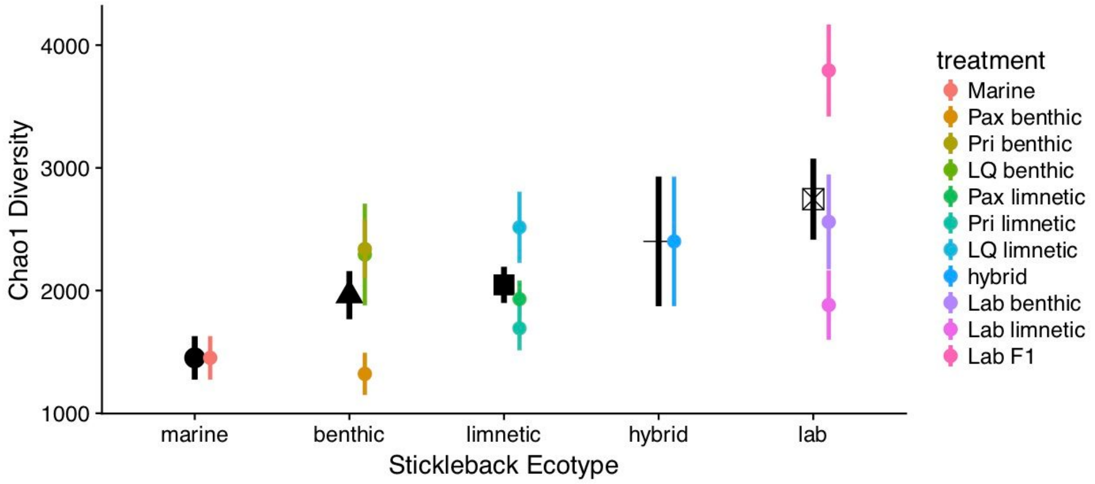

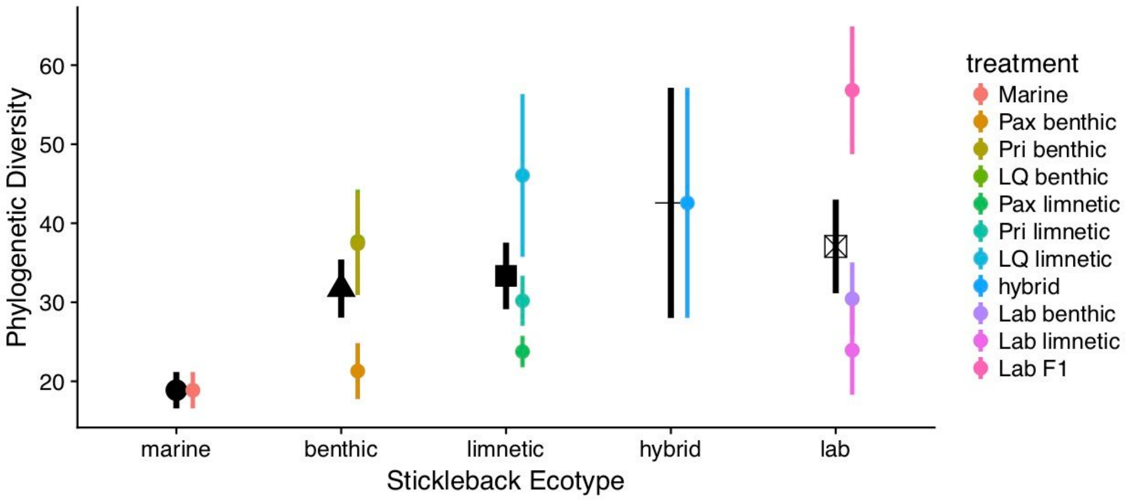

Phylogenetic Diversity
60
50
40
30
20
marine
benthic
limnetic
hybrid
lab
Stickleback Ecotype
treatment
Marine
Pax benthic
Pri benthic
LQ benthic
Pax limnetic
Pri limnetic
LQ limnetic
hybrid
Lab benthic
Lab limnetic
Lab F1

Supplementary Figure 3:

Results of PICRUSt analysis showing relative abundance of KEGG orthologs for pure benthic and limnetic ecotypes relative to the hybrid swarm from Enos lake.

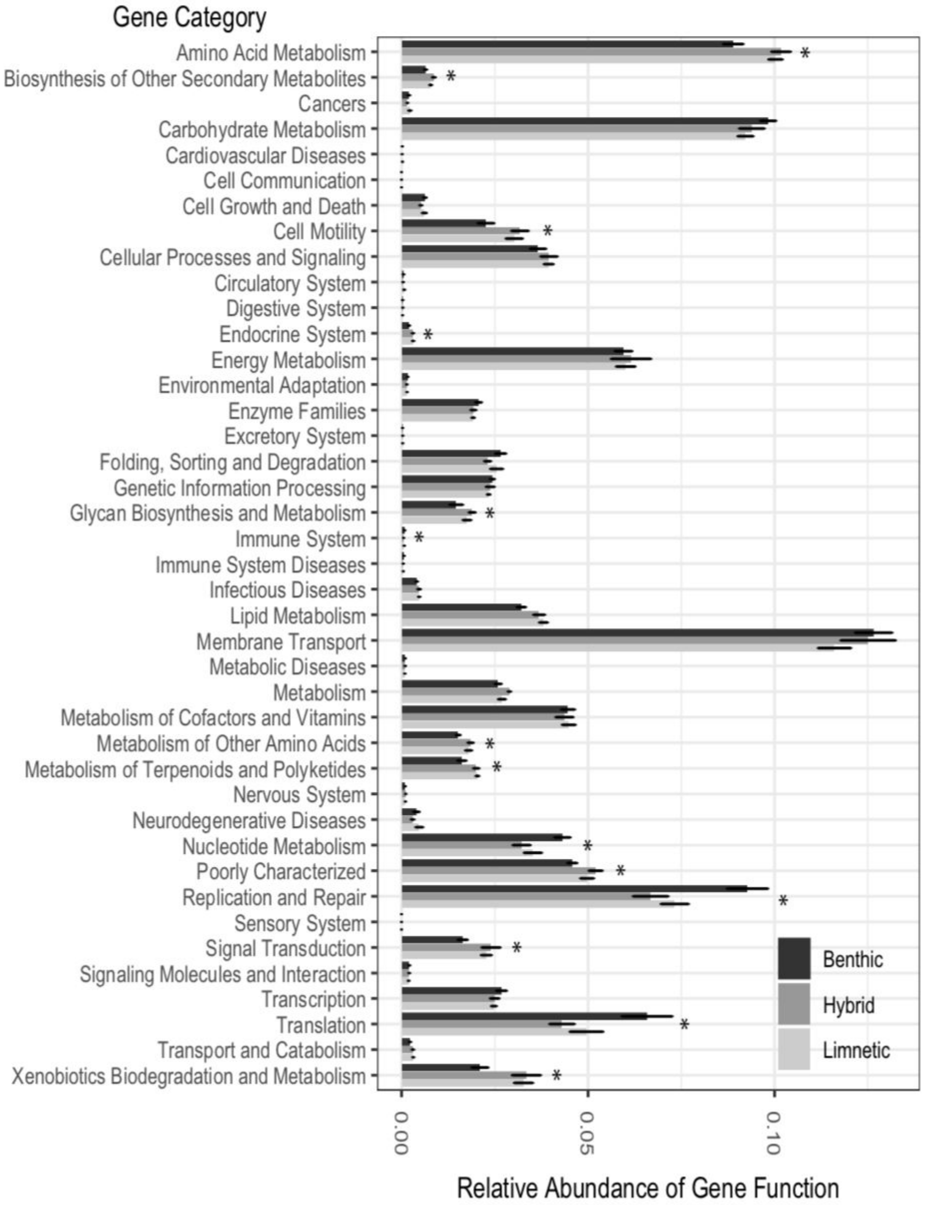

Supplementary Figure 4. Differentiation of the composition of the gut microbiome of lab reared benthic, limnetic, and F1 benthic-limnetic hybrid threespine stickleback. Multidimensional diversity is estimated using the Bray Curtis beta diversity metric.

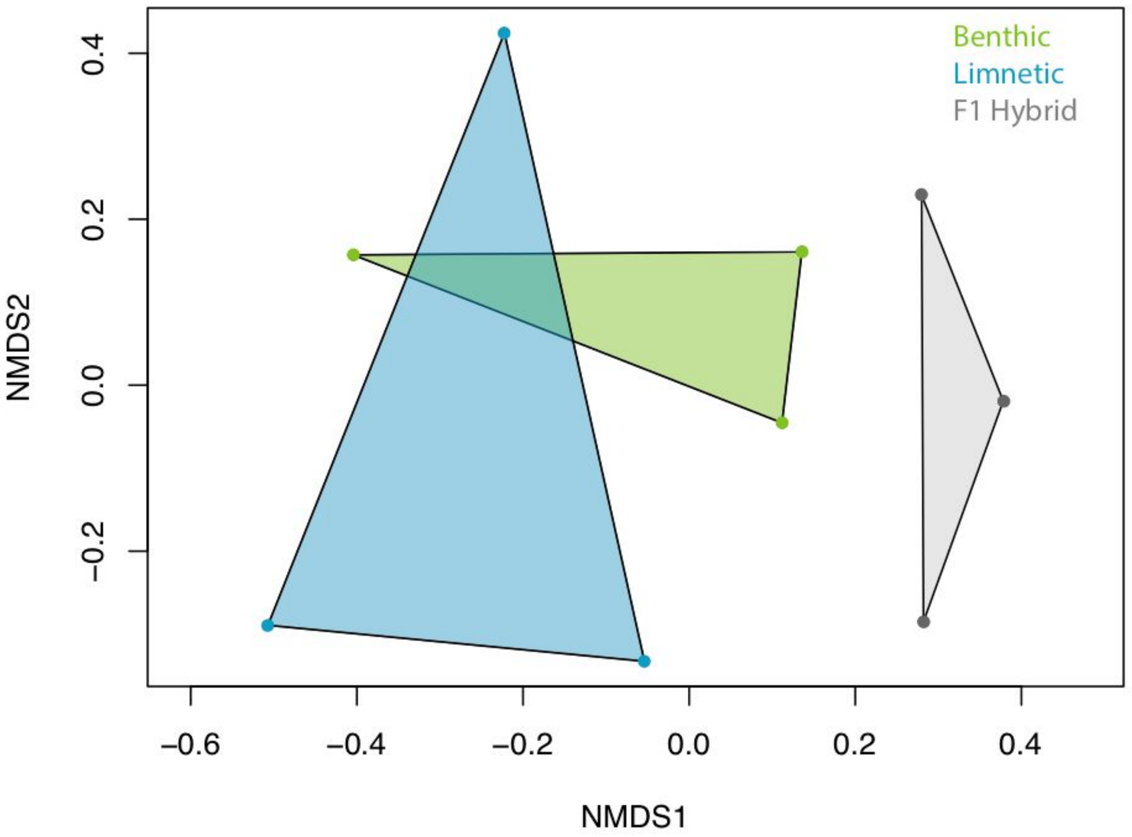
